# Mapping nonhuman cultures with the Animal Culture Database

**DOI:** 10.1101/2025.02.26.640400

**Authors:** Kiran Basava, Md Nafis Ul Alam, Liam Roberts, Kristen Martinet, Paige Cherry, Hector Garcia-Verdugo, Cristian Román-Palacios

## Abstract

Socially transmitted behaviors are widespread across the animal kingdom, yet there is a lack of comprehensive datasets documenting their distribution and ecological significance. Knowledge of animal behavioral traditions could be essential for understanding many species’ responses to anthropogenic disturbances and further enhancing conservation efforts. Here, we introduce the first open-access database that synthesizes data on animal cultural behaviors and traditions. The Animal Culture Database (ACDB) contains descriptions of 128 behaviors including forms of vocal communication, migration, predator defense, foraging practices, habitat alteration, play, mating displays, and other social behaviors for a sample of 61 species. In addition to offering an open-access resource for researchers, educators, and conservationists, the ACDB represents a step toward recognizing the role of social learning in animal populations.

## Background and summary

We present the first comparative, open-access database of animal culture, the Animal Culture Database (ACDB). The ACDB synthesizes data from studies of behavioral traditions and social learning in wild animal populations collected through a systematic literature review. It includes behaviors considered by primary researchers as population-specific due to factors other than ecological or genetic differences and transmitted via social learning. These behaviors are described and categorized into domains (such as communication, foraging, migration, and habitat alteration), and when available, the form of cultural transmission is noted and described (i.e. vertical, horizontal, and/or oblique transmission). We further summarize occurrence data, sampling periods, and taxonomic information on a sample of animal populations worldwide. In the long term, the ACDB can help answer questions related to behavioral flexibility and species’ resilience or vulnerability to anthropogenic disturbances. Because the explicit motivation of this database is to aid understanding of how socially learned behaviors interact with human-driven changes to the environment, it focuses on wild (rather than domestic or captive) populations and naturally-emerging behavioral traditions that may be relevant to their long-term fitness in the context of changing environments. To facilitate the integration between studies of animal culture and conservation measures, the ACDB documents effects on behaviors and populations from human activity when available from the literature. The ACDB is deployed as a web app at https://datadiversitylab.github.io/ACDB.

Decades of research have documented socially transmitted information forming what have been termed behavioral traditions or cultures in nonhuman animal populations worldwide^1–3^. These traditions include vocal dialects, migration routes, tool-use preferences, and foraging behaviors, which interact with species-typical social structures and behavioral flexibility^4^. Such behavioral diversity is a potentially neglected aspect of current conservation goals, as cognitive and behavioral flexibility have been shown to affect the ability of species to respond to environmental changes, disperse to different habitats, or adapt to human encroachment^5–7^. Despite calls to consider animal cultures when developing conservation strategies^8–10^, it is currently unclear how widely cultural behaviors are distributed globally and across the animal tree of life.

Comparative databases have increasingly been used in cultural evolutionary research on human societies to quantitatively test research questions that had previously been explored only qualitatively or in specific cultural settings^11^. While there is extensive literature on behavioral traditions and cultural transmission in a multitude of nonhuman species^3,12^, there is currently no centralized resource where quantitative data on animal culture can be accessed for analogous research objectives. The Animal Culture Database (ACDB) is an effort to comprehensively catalog socially transmitted behaviors across nonhuman animals. Many species display social learning, but only a proportion of these have recorded sets of behavioral repertoires in multiple domains^13,14^. Through assembling current scientific knowledge into a consolidated resource, the ACDB could help clarify when this variation is due to research bias and when it constitutes evidence for differences in reliance on cultural behaviors across taxa, highlighting geographic and taxonomic gaps where further primary research is needed. Additionally, the ACDB can be used to test associations between measures of cultural diversity and transmission and responses to anthropogenic disturbances.

A further goal of developing the ACDB is to encourage comparability of research across taxa, in part through integrating data highlighting unifying concepts of socially learned behaviors. For instance, a recent review of research by Rose et al.^15^ on bird vocal communication discussed the inconsistency in how researchers have historically defined vocalizations as ‘songs’ versus ‘calls’ and the need to instead focus on aspects of vocalizations along continuums^15^. The authors argue that this artificial distinction applies not only to the study of vocal communication within birds but across other taxa demonstrating vocal learning, including many mammalian species. Particularities like this may ultimately hinder comparative research on the evolution of vocal communication and social learning. Overall, we suggest that a consolidated resource could promote, when needed, the use of standardized frameworks for behavioral research across taxa.

### The culture concept

The Animal Culture Database includes published, primary observations of socially transmitted behaviors in wild populations. These observations also encompass work conducted in non-domesticated urban environments or semi-captive research settings. The existence, extent, and drivers of culture in nonhuman animals has been the subject of extensive discussion and debate over many decades^17–22^. For inclusion in the database, the criteria to which we have broadly adhered follows Boyd and Richerson^23^: “variation acquired and maintained by social learning”, or, what is called a “minimal definition of culture” by Schuppli and van Schaik^12^: “all behaviors and knowledge that are acquired and passed on within and between generations through social learning”. However, this describes what are considered only “traditions” by other researchers who, for instance, require multiple traditions for culture to be present^14,22^. There also remains debate over what constitutes sufficient evidence for the existence of social learning and culture depending on to what extent ecological, genetic, and other factors can be excluded as the causes of behavior.^12,20^

We have chosen a pragmatic and broadly inclusive approach for additions to the ACDB, using peer-reviewed studies that provide evidence of intraspecific behavioral variation between groups where social learning has not been excluded as a possibility by the primary researcher(s). It therefore remains for readers and users of the database to decide what they consider compelling evidence for culture in each population in the ACDB if they wish. To facilitate this, the ACDB includes a column in the Sources table that describes forms of evidence in each study. This column follows the taxonomy suggested by Whiten and Rutz^24^, a recent paper summarizing methodological approaches to identifying social learning across different animal culture studies. This is currently a qualitative, descriptive variable and will be standardized further in response to feedback on what is most useful to researchers.

The database at present is therefore agnostic with regard to a definition of animal culture. This was a pragmatic decision in line with its overall goal of facilitating conservation research. In their paper responding to Brakes et al.’s call^25^ for incorporating animal culture into conservation, Carvalho et al.^26^ point out the distinction between aiming to preserve behavioral/cultural diversity in a species for its own sake, or evaluating the conservation value of particular cultural behaviors. They caution against the use of the ‘culture concept’ in conservation in ways that bias towards charismatic groups that are already disproportionately the focus of research and conservation efforts. The same study also highlights the unlikeliness of reaching the evidence standards suggested by Brakes et al.^25^ such as demonstrating the effects of particular behaviors on survival and fitness, an attempt at which would slow existing conservation efforts. Even if intraspecific variation in behaviors between populations cannot be definitively established to be the result of social learning, we consider such variation worth recording and accounting for when attempting to preserve animal populations. Behavior is the result of continually interacting genetic, environmental and social factors, and even in humans, ecological and socially learned causes of cross-societal variation are difficult to disentangle^12,27^. We chose to sidestep these debates in our construction of this database with mind to its primary goal of aiding understanding of behavioral variation that may be relevant to conservation. Instead, we tend towards inclusion rather than exclusion of observed behaviors that could be socially learned and transmitted in wild populations to encompass groups that may not have been the focus of significant social learning research.

In addition to compiling research on animal culture, the ACDB further aims to enable researchers to examine the effects of climate change and human disturbances on cross-taxonomic patterns of animal behavior. Specifically, the ACDB currently includes a qualitative variable summarizing anthropogenic effects (discussed further in the Methods section). It also includes species’ IUCN Red List statuses as a proxy for extinction risk and standardized taxonomic data to facilitate interoperability with external datasets on environmental variability and anthropogenic stressors^28,29^. Ultimately, as the database expands to include more species and behaviors, data included in the ACDB could be used to explore questions related to environmental heterogeneity and cultural diversity, as well as associations between cultural diversity and conservation status in particular groups.

## Methods

We followed four major steps to construct the current release of the ACDB. First, we designed the database following relational database design principles^30^. Second, we conducted a systematic literature search to compile relevant published sources describing animal culture and related topics for particular groups. Third, we conducted a preliminary information extraction from the sources retrieved in the literature for main variables (e.g. culturally transmitted behaviors, behavioral categories, transmission modes, and population locations). Finally, we reviewed the initial coded data and transferred the resulting information into the relational database structure defined in the first step. Below, we provide additional details.

### Database structure

We followed principles of database normalization^31,32^ to structure the current release of the ACDB. Specifically, our relational database includes one subject per table with the goal of allowing 1) each species to be linked to multiple groups, 2) each group to be linked to multiple behaviors, and 3) multiple sources to be linked to each of the species, groups, and behaviors tables for different variables. The database includes a total of four tables. First, the species table contains taxonomic data for each species included in the database as well as species-specific information on social structure (Table 1). Second, the groups table contains data pertaining to each group of animals, including location, size, and where it falls in a species’ social structure when applicable (Table 2). This last item is particularly important in the case of species with multilevel social structure. One record in the species table can correspond to multiple records in the groups table. Third, the behaviors table contains descriptions for each cultural behavior recorded, including information on social transmission and potential effects from human activity (Table 3). One record in the groups table can correspond to multiple records in the behaviors table. Fourth, the sources table includes details on the relevant primary references for a given entry in the dataset (Table 4). Note that the species, groups, and behaviors table have fields specifying sources for particular variables (e.g. the primary social unit of a species, the size of a group, or the transmission mode of a behavior) linking them to the sources table. A comprehensive description of the database is shown in the diagram in Figure 2.

**Table 1.** Definitions for species-level variables.

| **species** |  |
| --- | --- |
| `species_id` [primary key] | Scientific name of the organism(s) in the format `Genus_species`. If the population/behavior was attributed to a subspecies, formatted as `Genus_species_epithet`. If the source(s) describing the group or behavior used an older version of the species name, the accepted form of the species name in GBIF was used. |
| `common_name` | Common name of the species in English |
| `GBIF` | GBIF usageKey for the species following gbif.org/species/ |
| `canonicalName` | Canonical name from GBIF API |
| `Genus ...Phylum` | Taxonomic names from GBIF backbone |
| `primary_social_unit`<br>`unit_evidence`<br>`unit_source` | The largest social unit for this species in which individuals maintain stable membership, adapted from Prox & Farine 2020 and Groot et al. 2023. Individual (no stable association with conspecifics); pair (two individuals, presumably mating pair); family (unit is composed of related individuals); group (unit includes unrelated individuals, may display fission-fusion behavior) |
| `IUCN` | Current IUCN Red List status |

**Table 2.** Definitions for variables on social groups within species.

| **groups** |  |
| --- | --- |
| `group_id` [primary key] | Unique ID for each group. Integer starting with 700000 |
| `species_id` | Foreign key linking to species table |
| `group_name` | Name of the group or population, e.g. Northern Resident killer whales or Gombe chimpanzees. Was assigned based on location if none appears to be commonly used. |
| group_level | The group level, relative to the species primary unit, to which cultural behaviors are attributed. 0+ with primary unit as 0; e.g. for killer whales 0 = matriline, 1 = pod, 2 = clan, 3 = community/population, 4 = ecotype, 5 = species |
| group_above | In the case of species with nested social structure, name of group directly above or encompassing the coded group; e.g. if focal group is Southern Resident killer whales, the group above this is the R-type or resident killer whale ecotype |
| size<br>size_evidence -<br>size_date -<br>size_source - | Approximate group size in number of individuals |
| location_evidence | As specific information possible on location, including research site, city, country, etc. |
| lat | Approximate latitude for location of population |
| long | Approximate longitude for location of population |

**Table 3.** Definitions of variables describing behaviors for groups.

| behaviors |  |
| --- | --- |
| group_id | Foreign key linking to groups table |
| behavior_id [primary key] | Unique ID for each behavior. Integer starting with 2. |
| behavior | Short name describing behavior, e.g. vocal dialect, termite fishing, lobtail feeding |
| behavior_description | Description of the behavior, placed in context of the species and the focal population(s) |
| behavior_source | Primary source used to describe the behavior |
| start_date | First observed date of behavior in the literature |
| end_date | Most recent observed date of behavior in the literature |
| vertical<br>vertical_evidence -<br>vertical_source - | Behavior is transmitted from parents to offspring |
| horizontal<br>horizontal_evidence -<br>horizontal_source - | Behavior is transmitted among peers in the same generation |
| oblique<br>oblique_evidence -<br>oblique_source | Behavior is transmitted from unrelated adults to juveniles |
| domains<br>domains_evidence -<br>domains_source | One of more of migration (knowledge of migration routes or sites); foraging (methods or tools of hunting or obtaining food, knowledge of food or resource patches, etc.); vocal communication (vocal transmission of information to others, e.g. songs and vocal dialects); antipredation (predator knowledge and avoidance); play (nonfunctional behavior that appears to be 'for fun'); mating (mating displays, breeding sites, mate selection); architecture (modifications to the physical environment- nest building, burrows, etc.); social (non-mating social behaviors, such as those that reinforce group identity) |
| anth_effects<br>anth_source - | Any effects of human activity on the behavior |

**Table 4.** Metadata for variable sources and references.

| sources |  |
| --- | --- |
| source_id [primary key] | Unique ID for each source in format firstauthoryear, e.g. baker2004 |
| title -<br>year -<br>authors -<br>doi - |  |
| methods and evidence | Description of the methods used to establish the presence of social learning and the existence of cultural behavior in the study population(s). |

**Figure 1.**
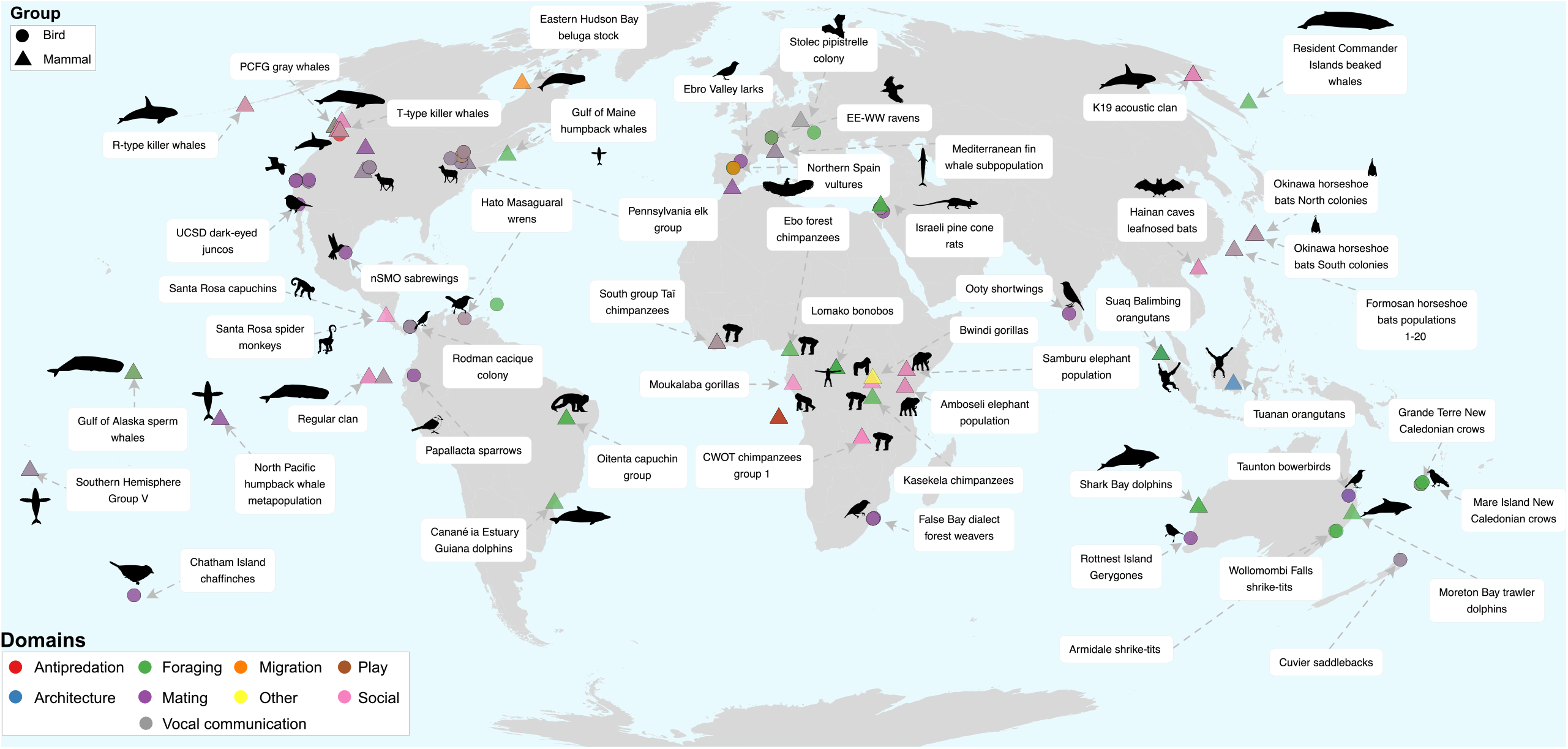
World map showing the geographic distribution of behavior records included in the current release of the Animal Culture Database. Colors indicate domains under which behaviors are classified. Shapes indicate taxonomic class (bird or mammal). A selected number of groups to which the behaviors are attributed are labeled. Images from phylopic.org^16^.

**Figure 2.**
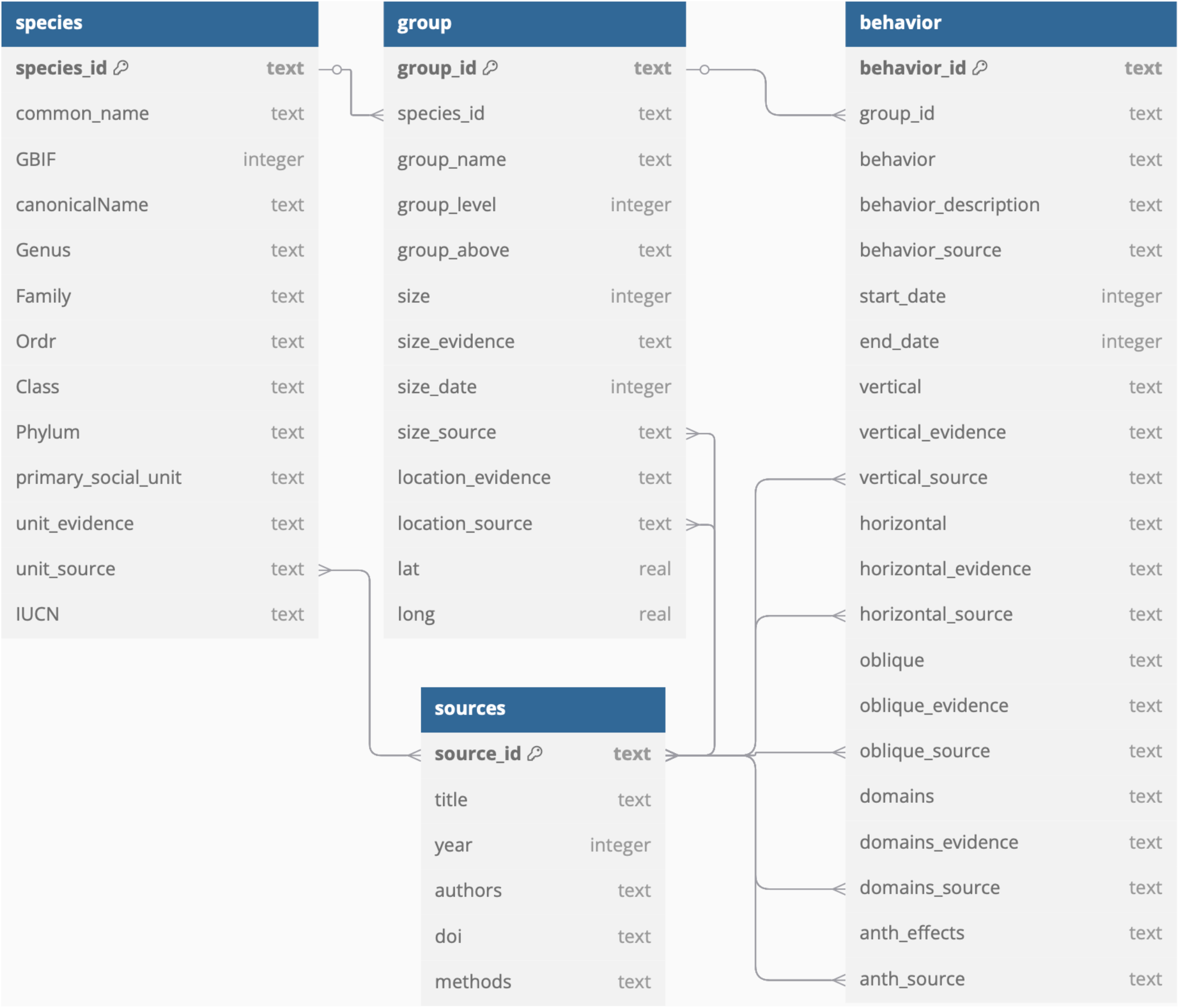
Structure of the relational database used to construct the Animal Culture Database. The four linked tables are shown with lines indicating one-to-many or many-to-many relationships. Key symbols indicate primary keys for each table.

### Literature search

We conducted a systematic literature review of published articles examining behavioral traditions and social learning across non-human animals in June 2024. We retrieved relevant papers using the Web of Science search engine based on terms in the title, abstract, and keywords, using the following search terms: (TS=(“animal culture” OR “behavio*ral tradition*” OR “cultural evolution” OR “cumulative culture” OR “cultural* transmi*” OR “cultural behavio*r” OR “cultural tradition*” OR “cultural drift” OR “vocal dialect*”)) AND TS=(“non-human animals” OR “nonhuman animals” OR “bird*” OR “aves” OR “avian” OR “primate*” OR “apes” OR “monkey*” OR “whale*” OR “insect*” OR “spider” OR “arachnid*” OR “arthropod*” OR “chiroptera*” OR “bats” OR “fish” OR “fishes” OR “mammal*” OR “ungulat*” OR “rodent*” OR “amphibia*” OR “reptil*”). These terms were chosen by reviewing papers on social learning and culture in different animal populations and noting frequently used phrases and terms (accounting for different spellings and phrasing), as well as terms for a wide range of taxonomic groups. This initial search resulted in a total of 5,490 papers. Once duplicates were removed using Zotero, the dataset consisted of 5,379 papers. Papers were then imported into Rayyan (rayyan.ai), an online tool for systematic reviews^33^ for an initial screening. Rayyan allows users to create lists of keywords for inclusion and exclusion that highlight terms and facilitate filtering of papers. Using Rayyan, we removed the most clearly irrelevant papers, including those on human behaviors, diseases, medical studies, and papers where animal culture referred to cell cultures. This first cleaning step in Rayyan reduced the original number of unique papers identified by the initial search from 5,379 to 2,364. We then manually curated the resulting dataset of articles through excluding conference abstracts, theses, reviews, and methodological papers and studies focused on non-animal groups. We restricted inclusion to papers containing original data on behavior in wild animal populations. This manual filtering resulted in a total of 1,058 papers, which formed the basis for the sources table in the relational database described above.

### Qualitative coding of animal culture

The initial coding of the database was focused on extracting qualitative descriptions of relevant behaviors from each paper using a modified schema from the final relational database structure. Papers with original data on animal behavior were tagged with a unique key in Zotero (generated by Rayyan) to link with the working dataset, and following data extraction tagged ‘read’. Papers which appeared to be relevant but not available were tagged ‘unavailable’, and those which appeared relevant but were not in English were tagged with the language to flag for later review; however, for the current release they were excluded. Papers which were not relevant (review papers or those not relating to animal behavior) were tagged ‘irrelevant’. For the relevant papers, we extracted taxonomic information, location, dates, and descriptions of possible culturally-transmitted behavior. Taxonomic names were matched to their records in the Global Biodiversity Information Facility (GBIF) Backbone Taxonomy^34^ to ensure consistency and interoperability with other datasets. The GBIF usageKey is included as a column in the species table containing taxonomic information as variable ‘GBIF’.

### Transfer to relational database

After the initial qualitative data extraction, the dataset was reviewed to determine variables for a standardized coding system. The dataset was then converted from CSV files to a relational database structure. This version of the database is stored as a SQLite database. The structure of the database was initialized using SQLiteStudio^35^ and was built in R using the packages DBI^36^ and RSQLite^37^. The relational structure of the database is shown in Figure 2. The variables for the species, groups, behaviors, and sources tables are defined in Tables 1-4 respectively and in the GitHub repository associated with this publication. In the tables, all variables named _evidence indicate free text descriptions or evidence for the above coded variable. All variables named _source indicate source for the above variable code and evidence, in the source_id format to link with the sources table.

## Data Records

The SQLite database and CSV files for the individual tables which form the database are available on figshare^38^ and in a GitHub repository (https://github.com/datadiversitylab/ACDB_datarelease). These are the files ACDB_v01.sql, and species.csv, groups.csv, behaviors.csv, and sources.csv. The R script used to construct the SQLite database from the CSV files is included as ACDB_sqlite_build.R. To view and interact with the database in R, the script example_queries.R is included in the repository with the required libraries and some example queries of the tables. Variables for each table are described in Tables 1-4 with the primary and foreign keys linking tables indicated. The web app associated with the ACDB can be found at: https://datadiversitylab.github.io/ACDB/.

## Technical Validation

For the construction of the initial qualitative dataset, papers were divided and screened by seven coders (KB, MD, LR, KM, PC, HGV, CRP). To facilitate standardization among coders, we held an initial collaborative training session where the lead author presented the coding approach and we discussed potential challenges and answered questions about the information extraction process. This session was also used to code an initial set of articles. We held weekly meetings for approximately five months to code papers and discuss potential problems in data extraction or solve general questions from coders. During transfer to the relational database, the leading author reviewed this dataset to assess data quality and fix potentially inaccurate codes by referring to the original papers and recoded the majority of papers and variables to adhere to standardized definitions (Table 1).

### The Animal Culture Database V1.0.0

The current release of the database contains data from a total of 121 papers on 30 mammal, 30 bird, and 1 insect species. Most of the records in the database are from groups in the tropics (i.e. temperate=40%, tropical=60%). Behaviors can be classified under one or more domains. Sixty-three behaviors were nonexclusively tagged as vocal communication, 44 as foraging, 39 as mating, 35 as social, five as architecture, four as migration and antipredation, two as play, and one as other (not fitting under any of the categories). As many of the behaviors were vocal displays used to attract mates (primarily song in birds and in some whales), 35 of the behaviors tagged as mating were also tagged as vocal communication. A distribution of the domains for which behaviors were tagged, with the counts for each domain corresponding to behavioral records for Mammalia, Aves, and Insecta is shown in Figure 3. Of the species sampled in the database, 37 have an IUCN status of Least Concern, four are Near Threatened, three are Vulnerable, eight are Endangered, four are Critically Endangered, and two are Data Deficient.

**Figure 3.**
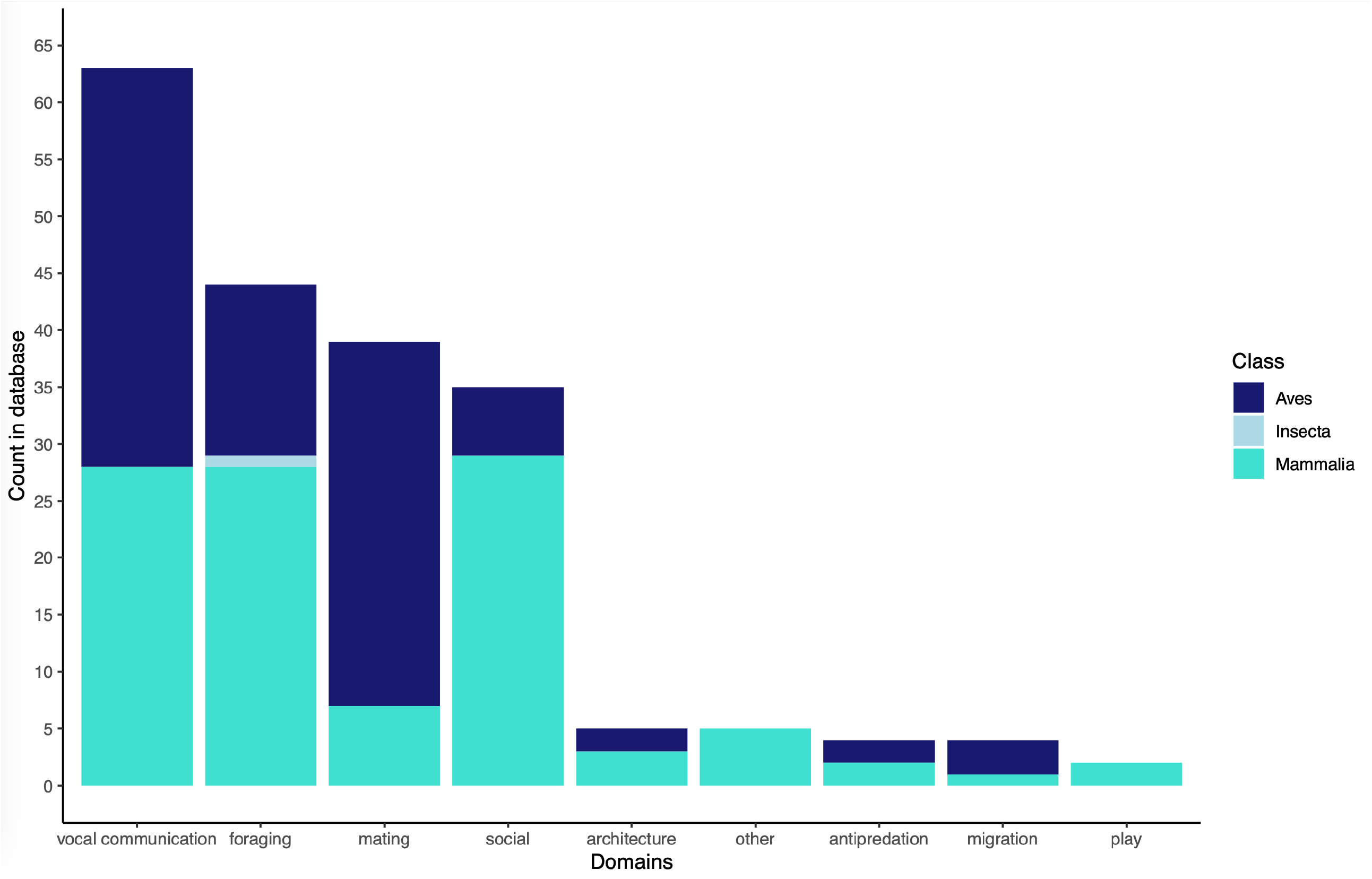
Count of domains under which behaviors are classified in the behaviors table, indicating counts for Aves, Mammalia, and Insecta.

### Limitations

The current release of the database encompasses only a sample of potentially relevant populations and behaviors. There are also clear biases to the data in the current version of the database. In particular, although the literature search used terms to encompass as broad a range of animal taxa as possible, sufficient research was only found on wild populations of mammal and bird species. Although there is evidence for social learning in other taxa, including invertebrate groups such as bees^30^, our search only retrieved experimental studies on these groups (e.g.^39^). The sample of taxa in the database broadly reflects overall research effort in animal behavior^40,41^. Analyzing the interaction between animal cultural behaviors and responses to anthropogenic disturbances while considering such discrepancies in research effort in conjunction with previously observed biases in conservation research^42^ will be an ongoing challenge. Addressing this can include ongoing testing of taxonomic and geographic bias as the database expands and variables from more literature are included.

The coding of the first version of the database was not conducted by taxonomic experts on the relevant groups. While the variables defined so far are relatively straightforward, this is a potential limitation on accuracy and depth of coverage. To mitigate this, we have adapted guidelines for best practices from the construction of comparative cultural databases in the social sciences^11^ such as making the data open access, inviting feedback/corrections, and including qualitative evidence and references for each code. We are also developing a pipeline for integrating external contributions from animal behavior researchers.

At this point, the database contains a small sample of the total documented cultural traditions in nonhuman animals. We highlight two caveats relevant to our literature search. First, it was not possible for this search to capture all relevant papers. Second, not all of the papers from our initial literature search have been included in the database as of time of publication-coding is ongoing and there are still >600 papers remaining from the search. The current version is the result of several months of a collaborative coding effort, which is still ongoing. The release of version 1.0.0 reflects our defining a formal versioning system for the ACDB as well as a natural checkpoint to incorporate feedback from users. We provide details on our approach to updating the database and accounting for feedback, including the already-reviewed papers as well as newly identified ones, in the section below.

### Roadmap, maintenance, and versioning for the ACDB

Future work on the ACDB will be done through database releases focused on including missing instances of animal culture, curating the existing ones, and implementing feedback from users on the overall structure of the database. Major releases (e.g. v1.0.0, v2.0.0, v3.0.0) will be based on additions of 100+ populations. Minor releases (e.g. v1.1.0, v1.2.0) will be based on additions of less than 10 populations. Changes to the contents of the database (e.g. inclusion of a missing paper, fixing taxonomy, adding locations) will be treated as patches (e.g. v1.1.1, v1.1.2). We aim for a major release every one to two years. The need for minor releases and patches will be assessed on a yearly basis.

Alongside the coding of papers from our literature search, we are incorporating behavioral records from major taxon-specific reviews of research on animal culture in order to ensure that studies considered significant by experts of culture in different taxa will be added to the database. We are actively seeking feedback on the overall structure of the database, the manner in which the groups and behaviors have been coded and data extracted from the sources gone through thus far, and changes that would help make it more useful for the animal culture and conservation research communities. Users can provide feedback by directly emailing the corresponding author, through opening a GitHub issue, or following the guidelines in the ACDB’s web app. Feedback can be in the form of the structure of the database, new instances of behaviors for existing populations or additional populations displaying culture, missing citations, and novel locations, among others.

## Code Availability

R version 4.3.2 and SQLiteStudio^35^ were used to build the relational database and populate tables with data from the CSV files. The R script and CSV files are available in the GitHub repository at https://github.com/datadiversitylab/ACDB_datarelease.

## Acknowledgements

We thank Andrea Thomer for advice on database construction. Mason Youngblood and Luca Hahn provided feedback on the conception of the project.

## Author contributions

K.B. conceived of the database. K.B. and C.R.P. wrote the draft. K.B., M.D., L.R., K.M., P.C., H.G.V., and C.R.P. coded the initial data from the literature search. K.B. curated the data for the linked tables and coded the relational database with supervision from C.R.P. All authors reviewed and edited the manuscript.

## References

1. Aplin, L. M. Culture and cultural evolution in birds: a review of the evidence. Animal Behaviour 147, 179–187 (2019). 10.1016/j.anbehav.2018.05.001.

2. Rendell, L. & Whitehead, H. Culture in whales and dolphins. Behavioral and Brain Sciences 24, 309–324 (2001). 10.1017/S0140525×0100396X.

3. Whiten, A. The burgeoning reach of animal culture. Science 372, eabe6514 (2021). 10.1126/science.abe6514.

4. Allen, J. A. Community through Culture: From Insects to Whales: How Social Learning and Culture Manifest across Diverse Animal Communities. BioEssays 41, 1900060 (2019). 10.1002/bies.201900060.

5. Berger-Tal, O. et al. Integrating animal behavior and conservation biology: a conceptual framework. Behavioral Ecology 22, 236–239 (2011). 10.1093/beheco/arq224.

6. Caro, T. & Sherman, P. W. Vanishing behaviors. Conservation Letters 5, 159–166 (2012). 10.1111/j.1755-263X.2012.00224.x.

7. Greggor, A. L. Animal Culture in Biodiversity Conservation. in The Oxford Handbook of Cultural Evolution (eds. Tehrani, J. J., Kendal, J. & Kendal, R.) (Oxford University Press, 2024).

8. Brakes, P. et al. Animal cultures matter for conservation. Science (2019) doi:10.1126/science.aaw3557.

9. Keith, S. A. & Bull, J. W. Animal culture impacts species’ capacity to realise climate-driven range shifts. Ecography 40, (2017). 10.1111/ecog.02481.

10. Whitehead, H., Ford, J. K. B. & Horn, A. G. Using culturally transmitted behavior to help delineate conservation units for species at risk. Biological Conservation 285, 110239 (2023). 10.1016/j.biocon.2023.110239.

11. Slingerland, E. et al. Coding culture: challenges and recommendations for comparative cultural databases. Evolutionary Human Sciences 2, (2020). https://doi.org/1017/ehs.2020.30.

12. Schuppli, C. & Schaik, C. P. van. Animal cultures: how we’ve only seen the tip of the iceberg. Evolutionary Human Sciences 1, e2 (2019). 10.1017/ehs.2019.1.

13. Galef, B. G. & Laland, K. N. Social Learning in Animals: Empirical Studies and Theoretical Models. BioScience 55, 489 (2005). 10.1641/0006-3568(2005)055[0489:SLIAES]2.0.CO;2.

14. Whiten, A. Cultural Evolution in Animals. Annu. Rev. Ecol. Evol. Syst. 50, 27–48 (2019). https://doi.org/46/annurev-ecolsys-110218-025040

15. Rose, E. M., Prior, N. H. & Ball, G. F. The singing question: re-conceptualizing birdsong. Biological Reviews 97, 326–342 (2022). https://doi.org/1111/brv.12800.

16. Keesey, T. M. Permalink - PhyloPic. https://www.phylopic.org/permalinks/cf45ad18cb1f8ad844309a20fe6b69e03bfb219cff9c5b80a9ef3e9c481f4021 (2024).

17. Galef, B. G. The question of animal culture. Human nature (Hawthorne, N.Y.) 3, 157–78 (1992). 10.1007/BF02692251.

18. Kendal, R. L., Kendal, J. R., Hoppitt, W. & Laland, K. N. Identifying Social Learning in Animal Populations: A New ‘Option-Bias’ Method. PLOS ONE 4, (2009). 10.1371/journal.pone.0006541.

19. Laland, K. N. & Hoppitt, W. Do animals have culture? Evolutionary Anthropology 12, 150–159 (2003). 10.1002/evan.10111.

20. Laland, K. & Janik, V. The animal cultures debate. Trends in Ecology & Evolution 21, 542–547 (2006). 10.1016/j.tree.2006.06.005.

21. Tennie, C., Call, J. & Tomasello, M. Ratcheting up the ratchet: on the evolution of cumulative culture. Phil. Trans. R. Soc. B 364, 2405–2415 (2009). 10.1098/rstb.2009.0052.

22. Whiten, A. & Van Schaik, C. P. The evolution of animal ‘cultures’ and social intelligence. Phil. Trans. R. Soc. B 362, 603–620 (2007). 10.1098/rstb.2006.1998.

23. Boyd, R. & Richerson, P. J. Why culture is common, but cultural evolution is rare. Proceedings of the British Academy, 88; Evolution of social behaviour patterns in primates and man 77–93 (1996).

24. Whiten, A. & Rutz, C. The growing methodological toolkit for identifying and studying social learning and culture in non-human animals. Phil. Trans. R. Soc. B 380, (2025). 10.1098/rstb.2024.0140.

25. Brakes, P. et al. A deepening understanding of animal culture suggests lessons for conservation. Proc. R. Soc. B. 288, rspb.2020.2718, 20202718 (2021). 10.1098/rspb.2020.2718.

26. Carvalho, S. et al. Using nonhuman culture in conservation requires careful and concerted action. Conservation Letters 15, (2022). https://doi.org/1111/conl.12860.

27. Mace, R. & Jordan, F. M. Macro-evolutionary studies of cultural diversity: a review of empirical studies of cultural transmission and cultural adaptation. Phil. Trans. R. Soc. B 366, 402–411 (2011). 10.1098/rstb.2010.0238.

28. Halpern, B. S. et al. Recent pace of change in human impact on the world’s ocean. Sci Rep 9, 11609 (2019). https://doi.org/1038/s41598-019-47201-9.

29. Fick, S. E. & Hijmans, R. J. WorldClim 2: new 1-km spatial resolution climate surfaces for global land areas. Intl Journal of Climatology 37, 4302–4315 (2017). https://doi.org/1002/joc.5086.

30. Harrington, J. L. The Relational Data Model. in Relational Database Design and Implementation 89–105 (Elsevier, 2016). 10.1016/B978-0-12-804399-8.00005-3.

31. Wickham, H. Tidy Data. J. Stat. Soft. 59, (2014). 10.18637/jss.v059.i10.

32. Allesina, S. & Wilmes, M. Computing Skills for Biologists: A Toolbox. (Princeton University Press, Princeton Oxford, 2019).

33. Ouzzani, M., Hammady, H., Fedorowicz, Z. & Elmagarmid, A. Rayyan—a web and mobile app for systematic reviews. Syst Rev 5, 210 (2016). https://doi.org/1186/s13643-016-0384-4

34. GBIF Secretariat. GBIF Backbone Taxonomy. Preprint at 10.15468/39OMEI (2023).

35. Salawa, P. SQLiteStudio. (2024).

36. DBI: R Database Interface. (2024).

37. Müller, K. et al. RSQLite: SQLite Interface for R. (2024). https://github.com/pawelsalawa/sqlitestudio/.

38. Basava, K. et al. The Animal Culture Database (ACDB) v1.0.0 data and code. University of Arizona Research Data Repository (2025). 10.25422/AZU.DATA.28850996.

39. Muth, F. Bumblebees show capacity for behavioral traditions. Learning & Behavior (2023) 10.3758/s13420-023-00594-0.

40. Bonnet, X., Shine, R. & Lourdais, O. Taxonomic chauvinism. Trends in Ecology & Evolution 17, 1–3 (2002). https://doi.org/1016/S0169-5347(01)02381-3.

41. Doody, J. S., Burghardt, G. M. & Dinets, V. Breaking the Social–Non-social Dichotomy: A Role for Reptiles in Vertebrate Social Behavior Research? Ethology 119, 95–103 (2013). https://doi.org/1111/eth.12047.

42. Dos Santos, J. W. et al. Drivers of taxonomic bias in conservation research: a global analysis of terrestrial mammals. Animal Conservation 23, 679–688 (2020). https://doi.org/1111/acv.12.

